# Bacteria-derived peptidoglycan triggers an NF-κB dependent response in *Drosophila* gustatory neurons

**DOI:** 10.1101/2021.12.13.472456

**Authors:** Ambra Masuzzo, Gérard Manière, Yaël Grosjean, Léopold Kurz, Julien Royet

## Abstract

Probing the external world is essential for eukaryotes to distinguish beneficial from pathogenic microorganisms. If it is clear that this task falls to the immune cells, recent work shows that neurons can also detect microbes, although the molecules and mechanisms involved are less characterized. In *Drosophila*, detection of bacteria-derived peptidoglycan by pattern recognition receptor (PRR) of the PGRP family expressed in immune cells, triggers NF-κB/IMD dependent signaling. We show here that one PGRP protein, called PGRP-LB, is expressed in some proboscis’s bitter taste neurons. *In vivo* calcium imaging reveals that the PGRP/IMD pathway is cell-autonomously required in these neurons to transduce the PGN signal. We finally show that NF-κB/IMD pathway activation in bitter neurons influences fly behavior. This demonstrates that flies use the same bacterial elicitor and signaling module to sense bacterial presence via the peripheral nervous system and trigger an anti-bacterial response in immune-competent cells.

## Introduction

Since microorganisms can reduce the fitness of their hosts, natural selection has favored defense mechanisms that protect them against disease-causing agents. The molecular mechanisms that are activated during the humoral and cellular responses, the main armed branches of the host against invading microbes, are known in great detail. However, to avoid a costly and not always efficient immune response, hosts can firstly engage in behaviors to avoid pathogenic microorganisms. Alternatively, they can adapt *adhoc* behaviors to reduce the consequences of the infection on themselves or their progeny. For instance, social insects, such as termites can ascertain the virulence of the *Metarhizium* and *Beauveria* fungi and avoid the most virulent strains^1^, while *Apis mellifera* workers are able to detect larvae infected with the fungus *Ascosphaera apis* and remove them from the nest^2^. On the other hand, since some microorganisms are beneficial for their host, animals can also be attracted by them. Up to date, the molecular and neuronal basis of these behavioral responses to microbes are much less characterized than the “canonical” immune responses. Genetically tractable models such as *Caenorhabditis elegans* or *Drosophila melanogaster* are very well suited to elucidate them ^3, 4, 5^.

Devoid of adaptative immunity like all invertebrates, *Drosophila* has emerged as a well-adapted model to unravel the signaling modules that control the innate immune responses against bacteria ^6, 7, 8, 9^. Essential to them are two NF-κB signaling pathways called Toll and IMmune Deficiency (IMD) whose activation triggers the production of immune effectors, such as AntiMicrobial Peptides (AMPs), in immune-competent cells ^6, 10, 11, 12^. This activation depends on the previous detection of bacteria-derived PeptidoGlycaN (PGN) by host Pattern Recognition Receptors (PRRs) belonging to the PeptidoGlycan Recognition Protein (PGRP) family ^13, 14^. Recent work has shown that signaling components of the NF-κB/IMD pathway, including the transcription factor Relish, and the upstream PGRP sensors are functionally required outside the immune system and more specifically in some neurons of the Central Nervous System (CNS) ^15^. Direct recognition of circulating bacteria-derived PGN by few brain octopaminergic neurons leads to their inhibition and, in turn, to an egg-laying reduction in PGN-exposed females ^16^. Hence, by detecting a ubiquitous bacteria cell wall component via dedicated PRRs, few brain neurons can adapt the female physiology to its infectious status.

The Peripheral Nervous System (PNS) of *Drosophila* and more specifically its gustatory and olfactory systems are also involved in microbe-induced behaviors. By activating a subclass of olfactory neurons that express the olfactory receptor Or56a, the microbial odorant Geosmin induces pathogen avoidance by inhibiting oviposition, chemotaxis, and feeding ^17^. In contrast, bacterial volatiles commonly produced during decomposition of plant material such as ammonia and certain amines, are highly attractive to flies ^18^. Furthermore, Or30a-dependent detection of bacteria-derived short-chain fatty acid induces attraction in larvae^19^. Previous works demonstrated that bacterial cell wall components like LipoPolySaccharide (LPS) and PGN are detected by *Drosophila’s* gustatory sensory system ^20^. Detection of LPS by the esophageal bitter Gustatory Receptor Neurons (GRNs) expressing the chemosensory cation channel TrpA1 (Transient receptor potential cation channel subfamily A member 1) triggers feeding and oviposition avoidance ^21^. PGN detection, instead, triggers grooming behavior upon stimulation of wing margins and legs but the nature of gustatory sensory neurons and receptors involved in this behavior remain elusive^22^.

We now present data demonstrating that the peptidoglycan interacting molecule PGRP-LB^23, 24, 25, 26^ is expressed in some GRNs housed in sensilla present on the fly labellum, a major organ of the peripheral taste system. We characterized PGRP-LB expressing GRNs and found that all of them belong to the labellar bitter neurons’ subclass. By performing calcium imaging, we revealed that bitter sensory neurons respond to PGN and that this response cell autonomously requires upstream elements of the NF-κB/IMD signaling cascade. Furthermore, we showed that manipulation of the NF-κB/IMD pathway activity in bitter GRNs leads to the modulation of feeding preference in a two choices-assay suggesting that PGN serve as a clue for flies to evaluate food sources. Moreover, our data indicate that NF-κB/IMD pathway activity in bitter GRNs modulates egg-laying behavior in females. These findings suggest that bacterial PGN elicits context-dependent host behaviors that are mediated by activation of the PGRP/IMD pathway in bitter neurons present in the proboscis. Combined with previous studies, these data demonstrate that the PGRP/IMD module is not only active in immune cells to trigger the production of antibacterial effectors but also in neurons of the PNS and the CNS to modulate the behavior of flies that are either in contact with or infected by bacteria.

## Results

### A peptidoglycan binding protein is expressed in some bitter gustatory neurons

Our previous work has shown that some PGN sensing molecules (PGRPs) are active outside immune cells and specifically in neurons of the CNS. Indeed, the direct detection of bacteria-derived PGN by the cytosolic protein PGRP-LE in some brain octopaminergic neurons modulates oviposition of infected females in an NF-κB-dependent manner ^15, 16^. To identify neurons which potentially expressed PGRPs and thus respond to PGN, we previously made use of a reporter line, pLB1^Gal4^, that partially recapitulates the endogenous expression of one PGRP protein (called PGRP-LB)^16^. We now noticed that, in addition to being expressed in some neurons of the brain, this line also labeled neuronal projections that originated from cells of the PNS. In pLB1^Gal4^/UAS-mCD8-GFP flies, a GFP signal was observed in the Sub-Esophageal Zone (SEZ) of the central brain where GRNs send their axonal projections (Fig. 1a, b) ^27, 28^. Accordingly, some spare cell bodies present in the labella at the position of taste sensory neurons were detected (here called pLB1+ neurons; Fig. 1c).Double staining between pLB1^Gal4^/UAS-mCD8-GFP and Gr66a-RFP, which is specifically expressed in bitter gustatory neurons, revealed that all pLB1+ neurons are bitter (Gr66a+), although they only represent a sub-population of them (Fig. 1d, e). Indeed, while there are around 25 Gr66a+ neurons per each labellum, we identified an average of 5 ± 2 pLB1+ neurons ^29, 30, 31^. We confirmed this result by using genetic intersectional strategy between pLB1^Gal4^ and Gr66a^LexA^ (Extended data Fig. 1a) and by using another driver that broadly targets bitter neurons (i.e., Gr32a^LexA^) (Extended data Fig. 1c). Consistently, by using the same strategy and a driver to label sweet GRNs (Gr5a^LexA^), we did not detect any neurons that are simultaneously pLB1+ and Gr5a+ (Extended data Fig. 1d). In addition, we assessed whether the expression of the Gal4 inhibitor Gal80 in Gr66a+ neurons (Gr66a^LexA^/LexAop^Gal80^) would suppress the expression of GFP in pLB1+ neurons (pLB1^Gal4^/UAS-mCD8-GFP). No signal was detected in pLB1^Gal4^/ UAS-mCD8-GFP flies expressing the Gal80 repressor, demonstrating that all the pLB1+ neurons in the proboscis are bitter (Extended data Fig. 1b). Lastly, imaging using a reporter line in which the endogenous PGRP-LB protein has been GFP-tagged at the locus (PGRP-LB::GFP) demonstrated that the endogenous PGRP-LB protein is also expressed in these neurons (Extended data Fig. 1e).

**Fig. 1.**
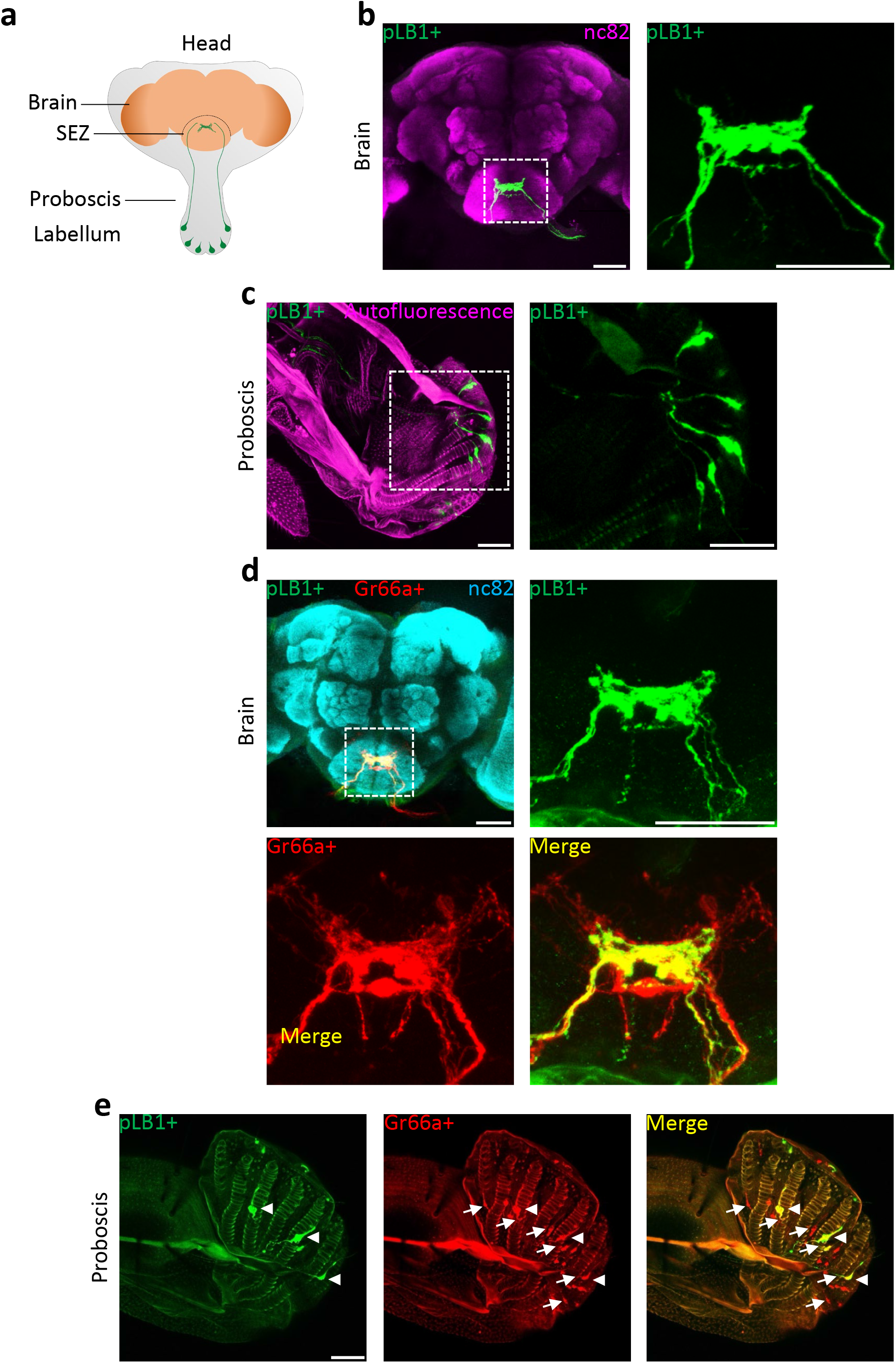
An IMD pathway component is expressed in neurons located in the proboscis. Immunodetection of cells expressing pLB1^Gal4^/UAS-mCD8-GFP (pLB1+). **a,** Schematic representing the fly head and the projections of pLB1+ peripheral neurons (green). The proboscis is an appendix dedicated to the feeding process and hosting neurons for the detection of tastants. The cell bodies of pLB1+ neurons are located in sensilla exposed to the environment and project axons to the brain, specifically in the sub-esophageal zone (SEZ). **b**, In the brain, pLB1+ cells send projections in the SEZ with a reproducible pattern. The panel on the right is a magnification of the SEZ delineated by the white box. **c**, The projections seen in the SEZ arise from neurons whose cell bodies are located in the tip of the proboscis, the labellum (sagittal view). The panel on the right is a magnification of the tip of the proboscis delineated by the white box. **d**, **e**, Immunodetection in the brain (**d**) and immunodetection in the proboscis (**e**) of cells expressing pLB1^Gal4^/UAS-mCD8-GFP (pLB1+) as well as Gr66a-RFP (Gr66a+). **d**, Top left is a view of a large portion of the brain, the other panels are magnifications of the sub-esophageal zone delineated by the white box. **e**, All the pLB1+ projections and cells (arrowheads) are Gr66a+ while not all the Gr66a+ projections and cells (arrows) are pLB1+. Scale bar, 50 μm.

### Bitter GRNs respond to DAP-type PGN

Two types of PGN, which differs for a single amino acid in the stem peptide, are found in bacteria. Whereas the Lysine (Lys)-type PGN is found in Gram-positive bacteria cell wall, the DiAminoPimelic acid (DAP)-type PGN forms that of Gram-negative bacteria. While Lys-type PGN preferentially triggers the *Drosophila* NF-κB/Toll pathway, DAP-type PGN mainly leads to the activation of the NF-κB/IMD pathway. By cleaving DAP-Type PGN in non-immunogenic fragments, PGRP-LB specifically modulates PGN levels and the activation of the NF-κB/IMD pathway. To test whether the pLB1+ cells were able to respond to PGN, we monitored their activity using the calcium sensor GCaMP6s. Exposing the labella of pLB1^Gal4^/UAS-GCaMP6s flies to DAP-type PGN, triggered an increase of the intracellular calcium levels in the axons of SEZ located pLB1+ neurons, indicating that those neurons sense and are activated by PGN (Fig. 2a, Extended data Movie 1). pLB1+ neurons responded to PGN in a dose-dependent manner and detected caffeine, but not sucrose, confirming their bitter nature (Fig. 2a, b, Extended data Movie 2). Considering that the pLB1^Gal4^ transgene drives the expression of Gal4 in neurons other than GRNs and in immune cells, and that all pLB1+ GRNs are GR66a+, we decided to study PGN perception by gustatory neurons in the well-characterized Gr66a+ GRN population. As for labellar pLB1+ neurons, calcium imaging revealed that DAP-type PGN activates Gr66a+ neurons (Fig. 2c, d, Extended data Movie 3) Together, these results showed that bitter GRNs, among which some express the PGRP-LB protein, are able to respond to DAP-type PGN. To evaluate the specificity of this response, pLB1^Gal4^/UAS-GCaMP6s and Gr66a^Gal4^/UAS-GCaMP6s flies were exposed to Lys-type PGN. When used at concentrations at which DAP-type PGN is active, Lys-type PGN was unable to trigger calcium increase in pLB1+, nor in GR66a+ neurons (Fig. 2e, f). These data indicate that bitter gustatory sensory neurons are specifically responsive to the PGN found in the cell wall of Gram-negative bacteria.

**Fig. 2.**
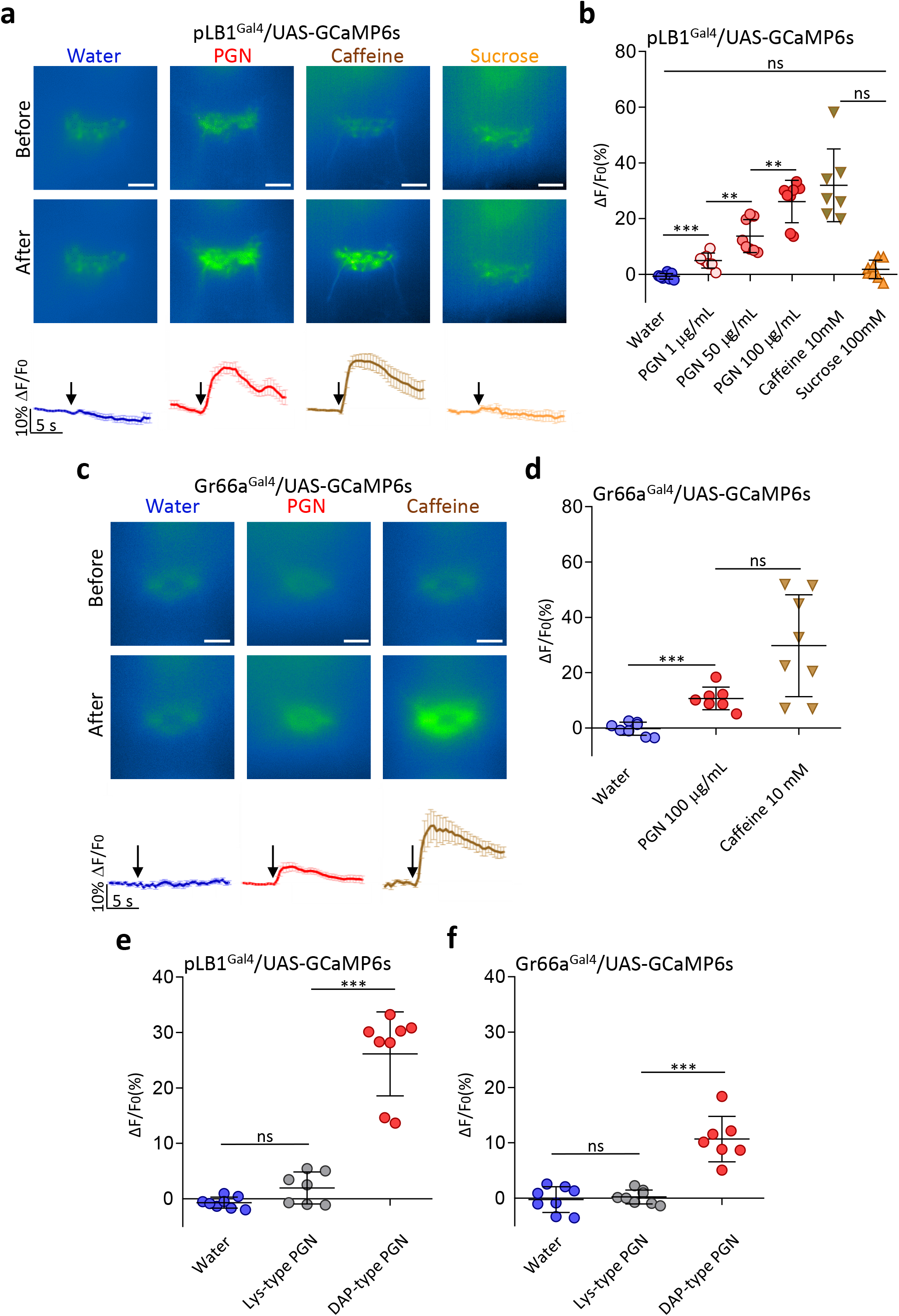
Bitter gustatory receptor neurons respond to DAP-type peptidoglycan. Real-time calcium imaging using the calcium indicator GCaMP6s to assess the *in vivo* neuronal activity in the sub-esophageal zone (SEZ) of pLB1+ neurons (pLB1^Gal4^/UAS-GCaMP6s) (**a, b**) or bitter gustatory receptor neurons (Gr66a^Gal4^/UAS-GCaMP6s) (**c, d**). **a** and **c**, Representative images (top) and averaged ± SEM time course of the GCaMP6s intensity variations (ΔF/F0%) (bottom). The addition of the chemical at a specific time is indicated by the arrow. The images illustrate the GCaMP6s intensity before and after the addition of either water as negative control (left panels), peptidoglycan (PGN; middle panels), caffeine or sucrose (right panels). The images represent the average intensity of 4 frames before or after the treatments. Scale bar, 20 μm. **b** and **d**, averaged fluorescence intensity of peaks ± SD for control, PGN, caffeine or sucrose -stimulated flies (N ≥7 for each treatment). **e** and **f**, Averaged fluorescence intensity of peaks ± SD for pLB1^Gal4^/UAS-GCaMP6s (**e**) or Gr66a^Gal4^/UAS-GCaMP6s (**f**) flies exposed to water, Lys-type PGN (100 μg/mL) or DAP-type PGN (100 μg/mL). *** indicates *p* < 0.0001; non-parametric t-test, Mann-Whitney test.

### Upstream elements of the NF-κB/IMD pathway are required for the response of bitter GRNs to PGN

Since some GRNs respond to Dap-type PGN, we tested whether the canonical upstream PGN sensors and downstream NF-κB/IMD pathway components were necessary to transduce the PGN signal, as it is for immune competent cells. For that purpose, *in vivo* calcium imaging experiments in pLB1^Gal4^/UAS-GCaMP6s flies were performed in various NF-κB/IMD mutant background flies. Two PGRP proteins function as upstream DAP-type PGN (hereafter simply PGN) receptors: PGRP-LC and PGRP-LE (Fig. 3a). While caffeine response was unaffected in *PGRP-LC* (*PGRP-LC^E12^*) and *PGRP-LE* (*PGRP-LE^112^*) mutants (Extended data Fig. 2a), PGN ability to activate pLB1+ neurons was completely abrogated in *PGRP-LC* mutants (Fig. 3b). In contrast, the response to PGN was weakly decreased in *PGRP-LE* animals. The loss of PGRP-LC was sufficient to abolish the response to PGN also in Gr66aGal4/UAS-GCamP6s flies, indicating that bitter gustatory neurons use mainly the membrane-associated receptor to detect the PGN (Fig. 3c). Since previous reports demonstrated that elements of the NF-κB/IMD pathway are expressed and functionally required in some neurons^15, 16^, their implication in mediating the effect of PGN was tested. While loss-of-function mutants for *Dredd* (*Dredd^D55^*) were responding normally to caffeine, a strong reduction of calcium signal was observed in flies stimulated with PGN (Fig. 3 b, c and Extended data Fig. 2a, c). The conserved ability of *Dredd* mutants to detect the caffeine demonstrated that their unresponsiveness to PGN was neither secondary to neuronal death nor to a loss of cell functionality. To ensure that the NF-κB/IMD pathway was required cell-autonomously in gustatory neurons, we used RNAi-mediated cell-specific inactivation. Functional downregulation of the PGRP-LC, IMD, Fadd, and Dredd in GR66a+ cells, was sufficient to block calcium response after PGN stimulation (Fig. 3d). These neurons remained responsive to caffeine (Extended data Fig.2d) demonstrating that the NF-κB/IMD pathway upstream components inactivation specifically impaired the response to PGN. Since most of the reported IMD-dependent responses have been shown to be mediated by the NF-κB transcription factor Relish, we tested its implication in bitter GRNs response to PGN ^6, 10^. Intriguingly, the calcium response of Gr66a+ neurons upon proboscis stimulation by PGN or caffeine was not statistically different in *Relish* RNAi flies compared to wild-type controls (Fig. 3d and Extended data Fig.2d). Altogether, these data demonstrate that Gr66a+ neurons can respond to DAP-type PGN in an IMD-pathway dependent manner, but independently of the canonical Relish trans-activator.

**Fig. 3.**
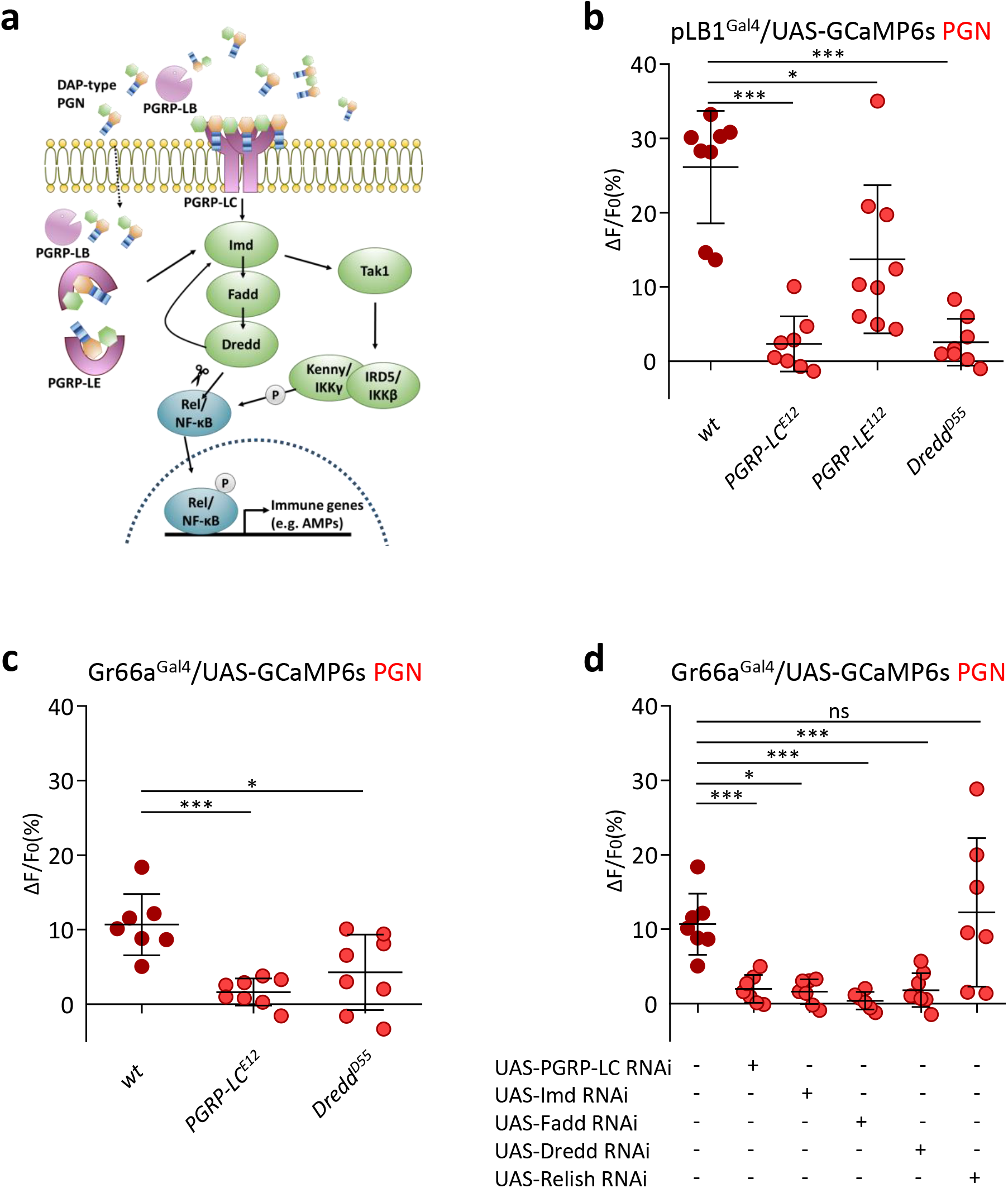
The PGN detection in pLB1+and Gr66a+ neurons require upstream elements of the NF-κB/IMD pathway. **a**, Schematic of the canonical NF-κB/IMD pathway in *Drosophila*. **b-d**, Real-time calcium imaging using the calcium indicator GCaMP6s to assess the *in vivo* neuronal activity in the sub-esophageal zone (SEZ) of pLB1+ neurons (pLB1^Gal4^/UAS-GCaMP6s) (**b**) or bitter gustatory receptor neurons (Gr66a^Gal4^/UAS-GCaMP6s) (**c,d**). **b,c**, Averaged fluorescence intensity of peaks ± SD for pLB1^Gal4^/UAS-GCaMP6s (**b**) or Gr66a^Gal4^/UAS-GCaMP6s (**c**) flies in different mutant backgrounds and exposed to PGN (100 μg/mL). **d**, Averaged fluorescence intensity of peaks ± SD for Gr66a^Gal4^/UAS-GCaMP6s animals expressing RNAi against different elements of the NF-κB/IMD pathway and exposed to PGN (100 μg/mL). **b-d**, 6≥N≥8 for each treatment. *** indicates *p*<0.0001; non-parametric t-test, Mann-Whitney test.

### The response of bitter neurons to peptidoglycan does not require TrpA1 nor Gr66a

A previous work has shown that another ubiquitous component of the Gram-negative bacterial cell wall, LPS, is detected in esophageal Gr66a+ bitter neurons via the TrpA1 cation channel ^21^. To assess whether TrpA1 is implicated in the response of pLB1+ neurons to PGN, we performed *in vivo* calcium imaging in *dTrpA1*mutants. The fact that PGN-dependent activation of pLB1+ cells is conserved in TrpA1 mutants demonstrated that PGN and LPS are detected by different receptors and certainly trigger different pathways in bitter GRNs (Extended data Fig. 2b). The non-GPCR gustatory receptor GR66a itself was also not involved in mediating the response to PGN. Altogether, these results suggest that PGRP-LC could be the dedicated receptor necessary for PGN detection and transduction in bitter neurons.

### Activation of the NF-κB/IMD pathway in bitter neurons modulate aversive behaviors

The ability of PGN to activate calcium release in bitter GRNs prompted us to test whether PGN triggers aversive behaviors in flies. We tested this hypothesis using the FlyPAD device in a two-choice feeding assay (Fig.4a)^32^. When flies were given a choice between a sucrose only and a sucrose plus PGN solution, no obvious repulsive behavior towards PGN was detected (Fig.4b and Extended data Fig. 3a, b). To further evaluate the phenotypical consequences associated with activation of the NF-κB/IMD pathway specifically in the Gr66a+ neurons, we overexpressed the upstream signaling receptor PGRP-LCa in these cells. This ectopic expression has been shown to induce forced dimer receptor formation and hence to trigger downstream signaling in the absence of the ligand or with lower amounts of it. In a two-choice feeding assay, flies in which PGRP-LCa was overexpressed in GR66a+ neurons, showed an increased repulsion towards solution containing PGN (Fig.4c). This behavior, which was not observed in control animals, was abolished by the simultaneous knockdown of the NF-κB/IMD downstream element Fadd (Fig. 4d). We then tested whether PGN-dependent activation of bitter neurons would impact egg-laying site preference. Although we were unable to detect any bias of egg-laying toward PGN contaminated media, we observed that PGRP-LCa overexpression in Gr66a+ neurons led to a decrease in oviposition (Fig. 4e, f). This decreased egg-laying when PGRP-LCa is expressed in bitter neurons was confirmed using Gr32a^Gal4^ as driver (Extended data Fig. 3c). These results suggesting that NF-κB/IMD pathway activation in bitter GRNs reduces female egg-laying were further confirmed by showing that this effect could be suppressed by the simultaneous RNAi-mediated Fadd inactivation in Gr66a+ cells (Fig. 4g). We previously demonstrated that PGN-dependent NF-κB/IMD pathway activation in a subset of brain octopaminergic neurons was sufficient to reduce female egg-laying^16^, a phenomenon reproduced with Kir2.1 overexpression in these cells, suggesting the PGN-dependent inactivation of the neurons. Importantly, inactivating the Gr66a+ cells via Kir2.1 expression did not phenocopy the egg-laying drop caused by inactivation of octopaminergic neurons suggesting that PGRP-LCa overexpression triggered activation of Gr66a+ neuron (Extended data Fig.3d). Consistently, conditional Gr66a+ cells activation via TrpA1 overexpression decreased female egg-laying (Fig. 4h). Taken together, these data demonstrate that receptor and transducers of the NF-κB/IMD pathway are expressed and functionally required in bitter neurons to mediate a behavioral response specifically towards DAP-type PGN.

**Fig. 4.**
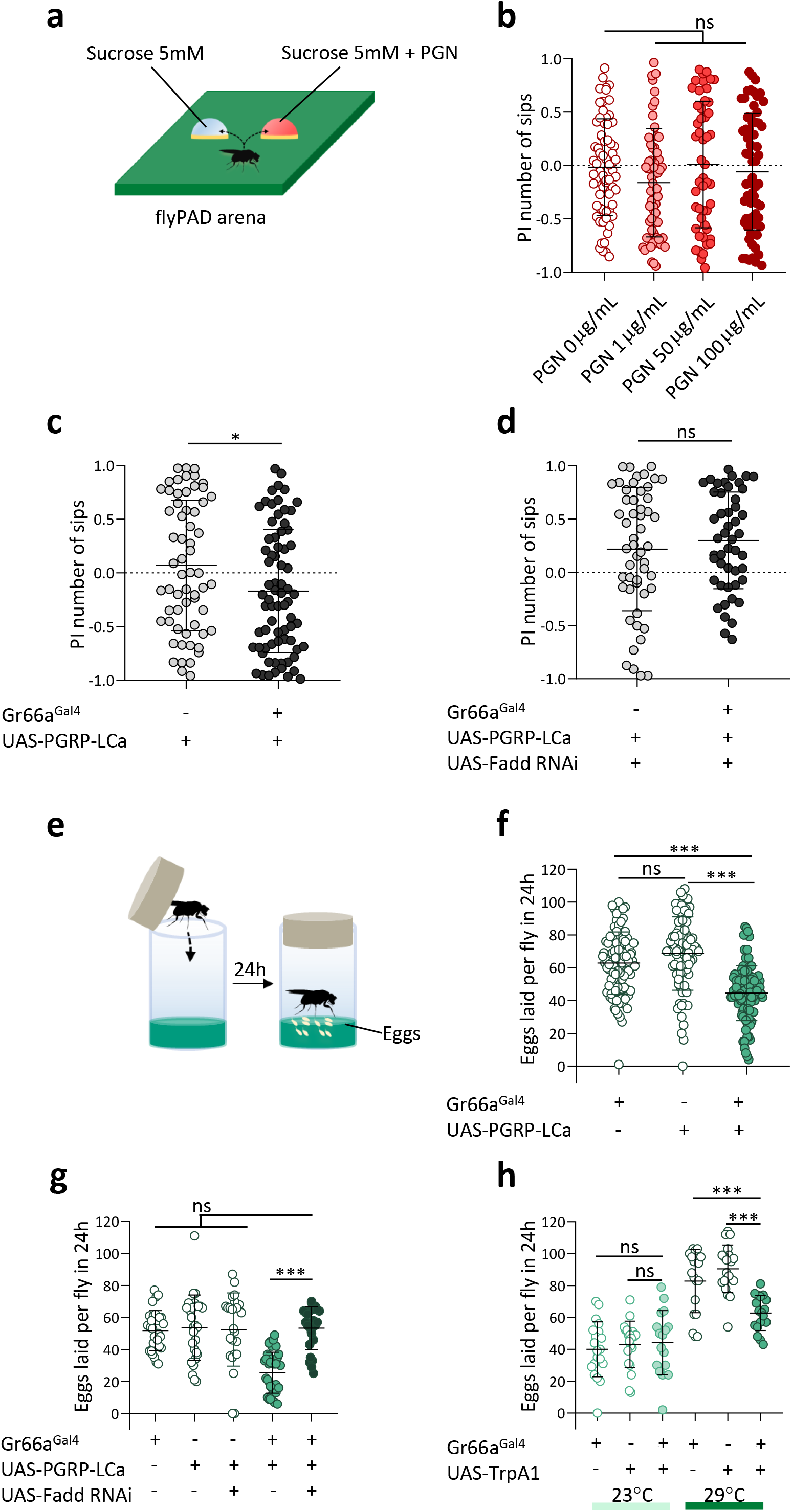
Activation of the NF-κB/IMD pathway activity in bitter neurons modulates feeding preference towards peptidoglycan and oviposition behavior. **a**, Schematic of the two-choice feeding assay using the flyPAD device. Individual flies are given the choice between a sucrose solution (5mM) and a sucrose solution (5mM) plus peptidoglycan (PGN) and tested for 1h. **b-d**, Feeding preference is expressed as a preference index (PI) based on the number of sips (see **Material and Methods**). **b**, Feeding preference of wild type (Canton S) flies exposed to two sucrose solutions (5mM), one of which containing PGN (different concentrations are tested and indicated in the X axis). **c**, Feeding preference of flies overexpressing PGRP-LCa in bitter taste neurons (Gr66a^Gal4^/UAS-PGRP-LCa) and controls exposed to two sucrose solutions (5mM), one of which containing PGN (100 μg/mL). **d**, Feeding preference of flies overexpressing simultaneously PGRP-LCa and UAS Fadd RNAi in bitter taste neurons (Gr66a^Gal4^/UAS-PGRP-LCa, UAS-Fadd RNAi) and control animals exposed to two sucrose solutions (5mM), one of which containing PGN (100 μg/mL). **e**, Schematic of oviposition assay. Individual flies are transferred in fresh tubes and allowed to lay eggs for 24h. **f**, Eggs laid per 24 hours (24h) by flies overexpressing PGRP-LCa in bitter taste neurons (Gr66a^Gal4^/UAS-PGRP-LCa) and control animals. **g**, Eggs laid per 24h by flies overexpressing simultaneously PGRP-LCa and Fadd RNAi in bitter neurons (Gr66^Gal4^/UAS-PGRP-LCa, UAS-Fadd RNAi) and control animals. **h**, Eggs laid per 24h by flies overexpressing TrpA1 in bitter neurons (Gr66^Gal4^/UAS-TrpA1) and control animals, at a permissive (23°C) and restrictive (29°C) temperature. **b-d**, shown are the average PI ± SD of at least three independent trials (N≥16 for each trial). *** indicates p<0.0001; ns indicates p > 0.05; non-parametric t-test, Mann-Whitney test. **f-h**, shown are the average numbers of eggs laid per fly per 24 h ± SD from at least two independent trials with at least 20 females per trial, genotype and condition used. *** indicates p<0.0001; ns indicates p > 0.05; non-parametric ANOVA, Dunn’s multiple comparison test.

## Discussion

This study demonstrates that some neurons of the gustatory system exposed to a microbe-contaminated environment are able to detect the presence of bacteria by sensing one of its main conserved and ubiquitous cell wall components, peptidoglycan. In bitter neurons, this detection is mediated by the IMD pathway PGRP-LC receptor and thus probably not by classical Gr proteins such as Gr66a. The PGN signal is transduced by the known cytosolic members of the IMD pathway such as Fadd and Dredd. Together with previous reports, these results confirm the key role played by the PGN/PGRP module in regulating many of the interactions between bacteria and flies. This specific recognition step, which takes place at the cell membrane via PGRP-LC or within the cells via PGRP-LE, has been shown to control the production of anti-bacterial effectors by immune-competent cells, to alter the egg-laying rate of infected females and to allow the physiological adaptation of the flies to their infectious status ^15, 16, 33, 34, 35, 36^. Interestingly however, while the initial MAMP/PRR recognition event is conserved among these processes, the downstream molecular mechanisms that transduce the signal are context-dependent. Whereas the PGN-dependent activation of an immune response in adipocytes, hemocytes or enterocytes and the inhibition of VUM III octopaminergic brain neurons rely on the nuclear NF-κB/Relish protein, the transcriptionally regulated effectors are likely to be different ^7, 16^. The AMPs that are the main NF-κB-dependent mediators of the antibacterial response, seem to be dispensable for the reduction of oviposition in infected flies (AM, LK, JR, personal communications). The response of bitter neurons to PGN uses a non-canonical IMD pathway in which NF-κB/Relish is not required. Interestingly, PGRP-LC and some downstream IMD components are also required at the pre-synaptic terminal of *Drosophila* motoneurons for robust presynaptic homeostatic plasticity ^37, 38^. The local modulation of the presynaptic vesicle release, which occurs in seconds following inhibition of postsynaptic glutamate receptors, required PGRP-LC, Tak1 but is also Relish-independent. These data and ours raise important questions regarding how the activation of the upstream elements of the IMD cascade is modifying neuronal activity, a topic for future studies. Previous biochemical studies have shown that IMD signaling is rapid, occurring in seconds, a time frame consistent with its role at the synapse and now in bitter neurons signal transduction^39^. Another possibility for the involvement of the IMD pathway in the bitter neurons would be that the expression of a yet to be identified PGN sensor requires the PGRP/IMD module for a permissive signal upon stimulation by environmental bacteria.

Our data show that flies can perceive PGN, a component of the bacteria cell wall, via bitter neurons triggering behaviors. These findings are complementary to observations made for another cell wall component in Gram-negative bacteria, called LPS, which triggers feeding avoidance in *Drosophila* through the activation of bitter neurons^21^. While LPS induced avoidance behavior is mediated through the chemosensory cation channel TrpA1, we show that PGN induced activation of bitter neurons seems to be independent of it. We demonstrate that the bitter response upon PGN stimulation is dependent on the IMD pathway that not only regulates a feeding aversion for PGN but also controls ovipositional avoidance. This indicates that PGN is probably an informative environmental cue for flies. Our approach is minimalist and allows us to directly link a molecule to the neurons that perceive it, but it is most probable that the behavior of flies in a natural environment corresponds to a highly complex integration of multiple intricate signals perceived by different sensory systems of the animal.

In nature, PGN is likely detected in combination with other tastants and odorants, which detected alone may lead to an array of conflicting behaviors but in combination will yield in one context-dependent behavioral output ^21, 40^. Consequently, it may be hazardous to expect clear phenotypes, or to make sense of the observed ones when testing a single molecule of the permanent environment of the fly while this molecule is not especially deleterious per se, but rather informative for the animal. The PGN is an interesting case as on one hand, an internal sensing of this molecule indicates an infection, the uncontrolled growth of a bacteria or a breach in a physical barrier. On the other hand, the perception of this same molecule in the environment might be a clue, among others, to suggest a heavily contaminated place. As far as PGN perception is concerned, other levels of regulation are expected ^17, 19, 41^. Indeed, the pLB1 line used in this study to probe the neurons that can respond to PGN is only labelling a subclass of GR66a positive neurons. This suggests that the response to PGN is likely to be not homogenous among all bitter neurons. Besides, the pLB1 line partially recapitulates the endogenous expression of PGRP-LB, an enzyme that by cleaving the PGN into non-immunogenic muropeptides, buffers the PGN-dependent IMD response ^34, 42, 43^. It is therefore possible that non-pLB1 bitter neurons will also respond to PGN and that this response might be attenuated in pLB1 neurons.

## Extended data

**Extended data Fig. 1.**
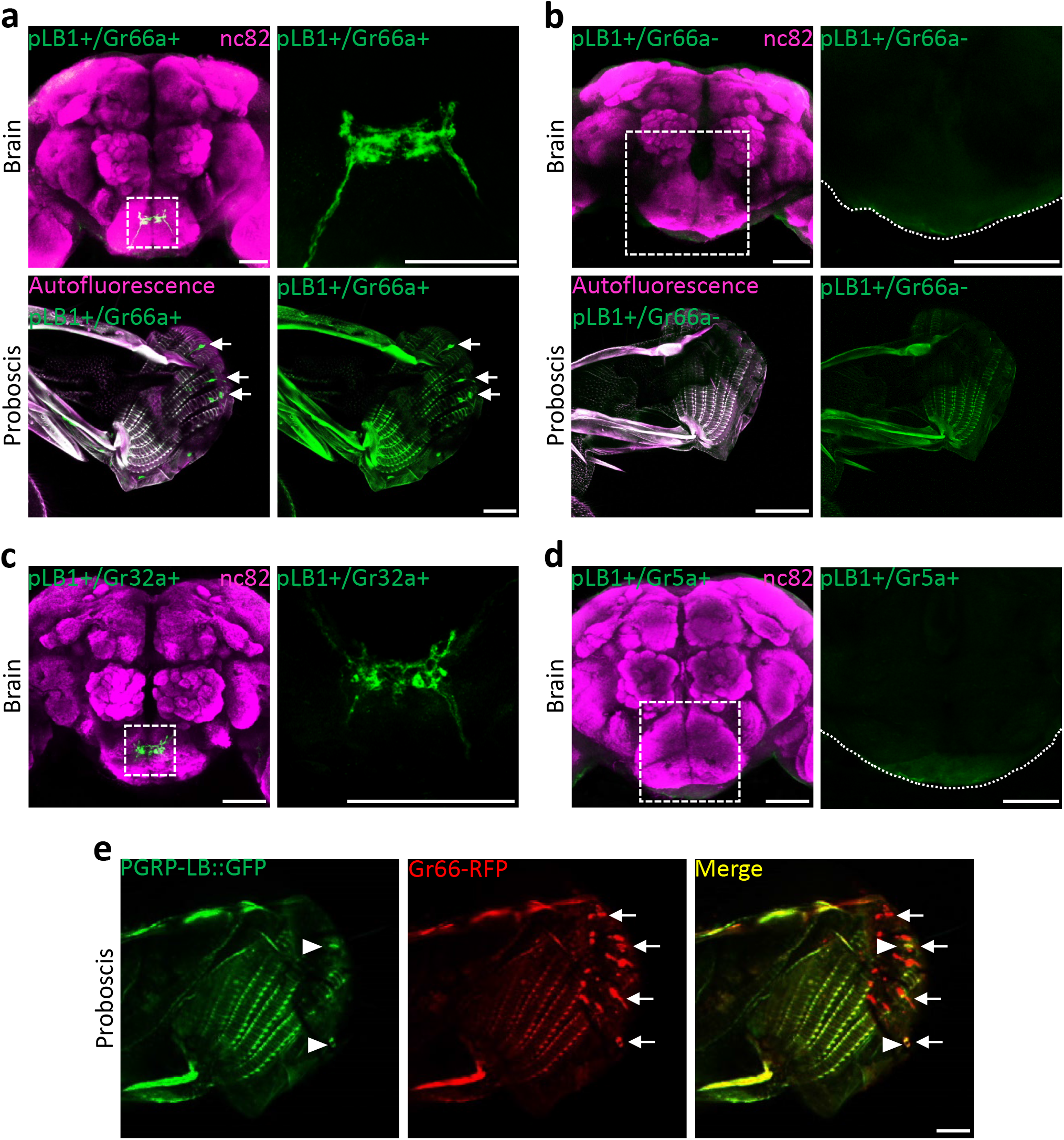
pLB1 neurons in the labellum are exclusively Gr66a. **a**, Immunodetection in brain (top) and immunodetection in the proboscis (bottom) of cells pLB1+ as well as Gr66a+ via genetic intersectional strategy (pLB1^Gal4^, Gr66a^LexA^/UAS*frt*STOP*frt*mCD8-GFP, LexAopFLP). Arrows point to pLB1+/Gr66a+ cellular bodies. **b**, Immunodetection in brain (top) and detection in the proboscis (bottom) of cells pLB1+ and Gr66a- (pLB1+/Gr66a-) via the expression of the Gal4 inhibitor Gal80 specifically in Gr66a+ cells (pLB1^Gal4^; UAS-mCD8-GFP/Gr66a^LexA^, LexAopGal80). **c**, Immunodetection in the brain of cells pLB1+ as well as Gr32a+ via genetic intersectional strategy (pLB1^Gal4^/Gr32a^LexA^; UAS*frt*STOP*frt*mCD8GFP, LexAopFLP). **d**, Immunodetection in the brain of cells pLB1+ as well as Gr5a+ via genetic intersectional strategy (pLB1^Gal4^, Gr5a^LexA^/UAS*frt*STOP*frt*mCD8GFP, LexAopFLP). **e**, Detection in the proboscis of cells producing the endogenous PGRP-LB (PGRP-LB::GFP) as well as Gr66a-RFP (Gr66a+). All the PGRP-LB::GFP+ cells (arrowheads) are Gr66a+ while not all the Gr66a+ cells (arrows) are PGRP-LB::GFP+. In **a-c**, the top right and right panels are magnifications of the sub-esophageal zone delineated by the white box. All the images of the proboscis are sagittal views. Scale bar, 50 μm.

**Extended data Fig. 2.**
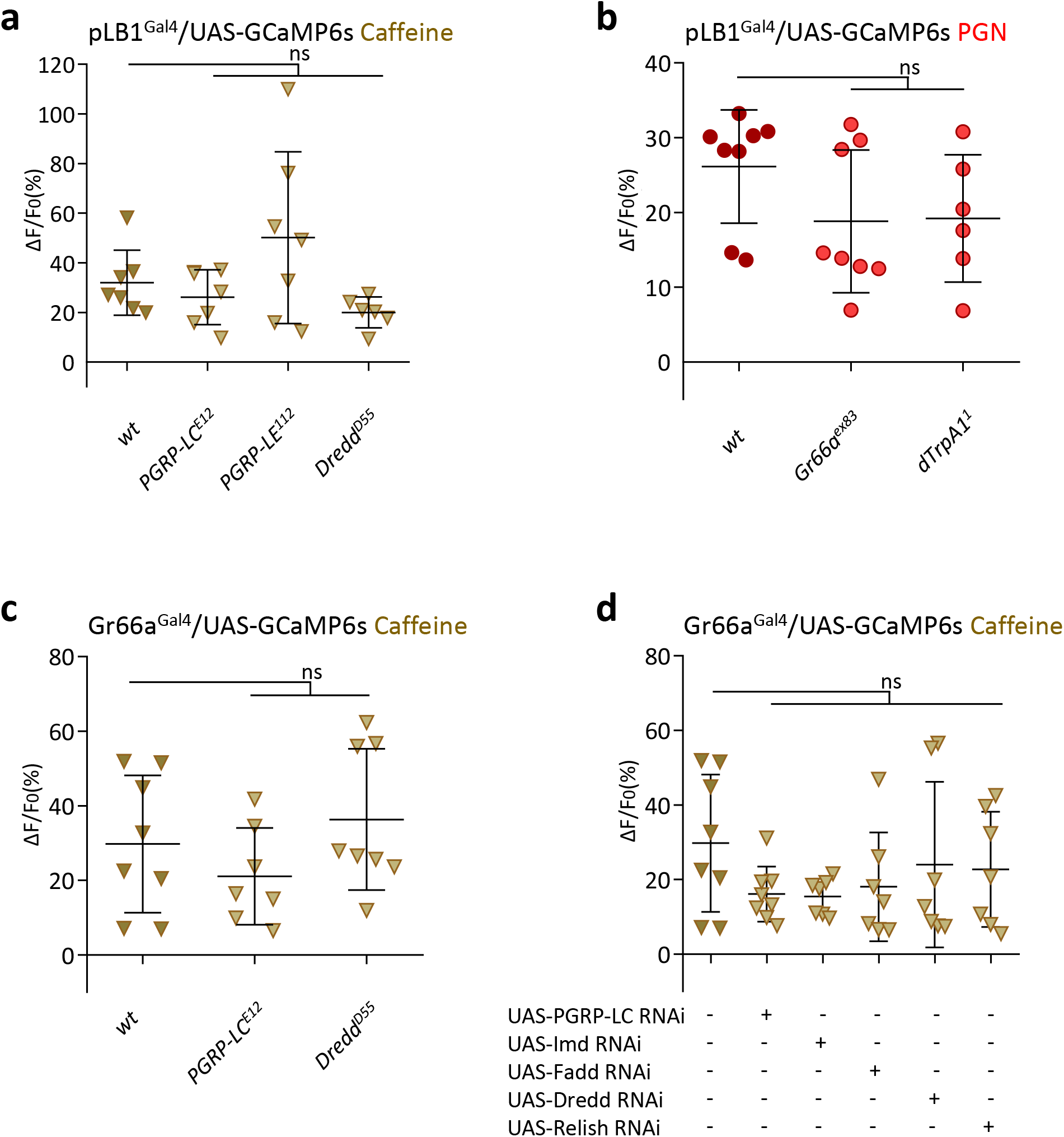
Bitter taste neuron response to caffeine. **pLB1+ neurons response to PGN does not require Gr66a or dTRPA1** Real-time calcium imaging using the calcium indicator GCaMP6s to assess the *in vivo* neuronal activity in the SEZ of pLB1+ (**a** and **b**) or Gr66a+ (**c** and **d**) neurons. **a,c**, Averaged fluorescence intensity of peaks ± SD for pLB1^Gal4^/UAS-GCaMP6s (**a**) or Gr66a^Gal4^/UAS-GCaMP6s (**c**) flies in different mutant backgrounds exposed to caffeine (10mM). **b**, Averaged fluorescence intensity of peaks ± SD for pLB1^Gal4^/UAS-GCaMP6s flies in different mutant backgrounds exposed to peptidoglycan (100 μg/mL). **d**, Averaged fluorescence intensity of peaks ± SD for Gr66a^Gal4^/UAS-GCaMP6s animals expressing RNAi against IMD pathway elements and exposed to caffeine (10mM). **a-d**, 6≥N≥8 for each condition. ns indicates *p* > 0.05; non-parametric t-test, Mann-Whitney test.

**Extended data Fig. 3.**
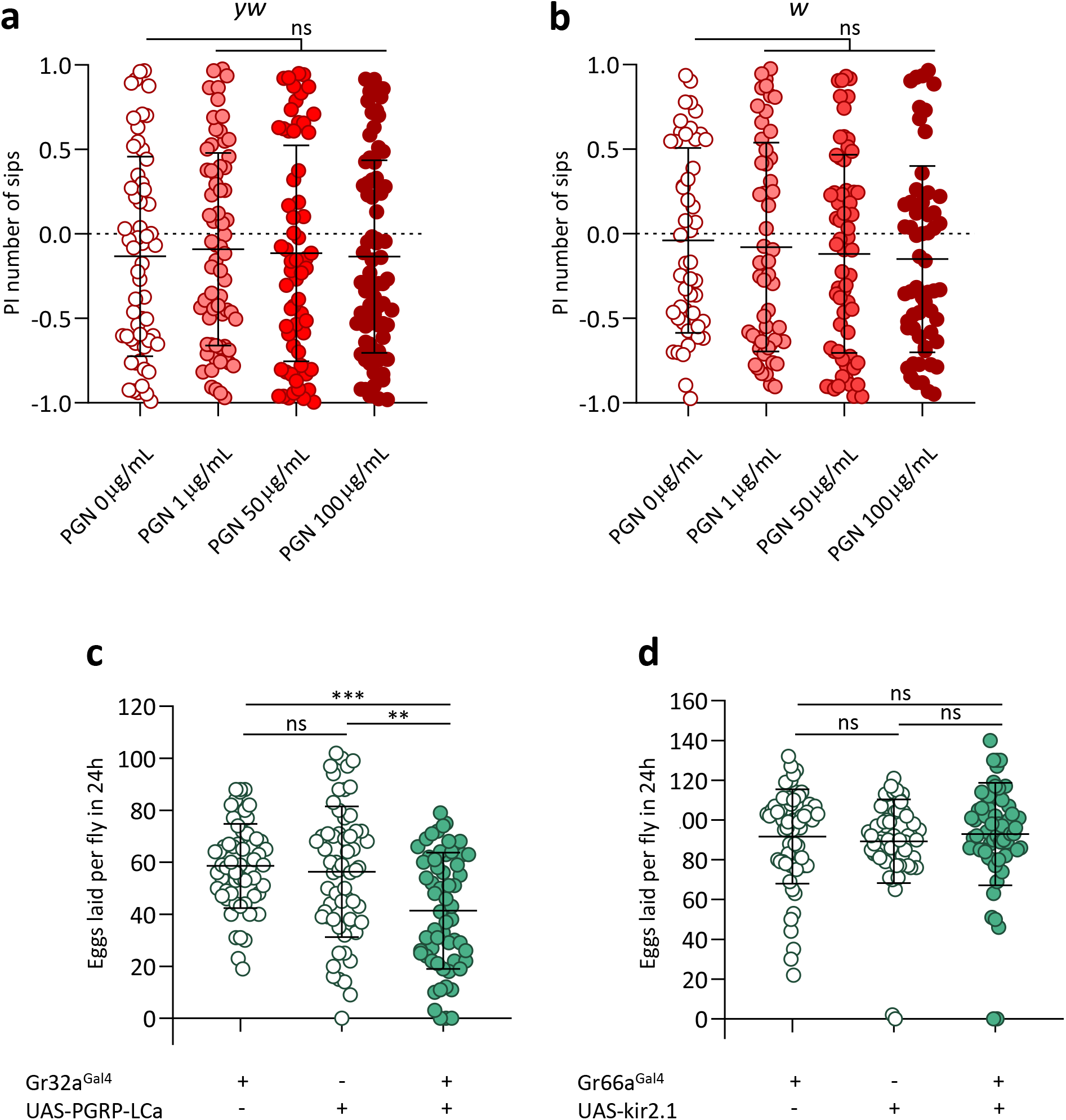
PGN is neither attractive nor aversive for wild type flies in two-choice feeding assay. IMD pathway activation in bitter neurons inhibits egg laying. **a,b**, Feeding preference of w (**a**) or yw (**b**) flies exposed to two sucrose solutions (5mM), one of which containing PGN (different concentrations are tested and indicated in the X axis). **c**, Eggs laid per 24 hours (24h) by flies overexpressing PGRP-LCa in bitter taste neurons (Gr32a^Gal4^/UAS-PGRP-LCa) and control animals. **d**, Eggs laid per 24 hours (24h) by flies overexpressing kir2.1 in bitter taste neurons (G66a^Gal4^/UAS-kir2.1) and control animals. **a,b**, Shown are the average PI ± SD of at least 5 independent trials (N≥16 for each trial). *** indicates p<0.0001; ns indicates p > 0.05; non-parametric t-test, Mann-Whitney test. **c,d**, shown are the average numbers of eggs laid per fly per 24 h ± SD from with at least 20 females per trial, genotype and condition used. *** indicates p<0.0001; ns indicates p > 0.05; non-parametric ANOVA, Dunn’s multiple comparison test.

**Extended data Movie 1 | pLB1+ neurons respond *in vivo* to PGN.**

Real-time calcium imaging using the calcium indicator GCaMP6s to assess the *in vivo* neuronal activity in the sub-esophageal zone of pLB1 neurons (pLB1^Gal4^/UAS-GCaMP6s). Effect of peptidoglycan solution stimulation (100 μg/mL). GFP signal was recorded every 500 ms.

**Extended data Movie 2 | pLB1+ neurons respond *in* vivo to caffeine.**

Real-time calcium imaging using the calcium indicator GCaMP6s to assess the *in vivo* neuronal activity in the sub-esophageal zone of pLB1 neurons (pLB1^Gal4^/UAS-GCaMP6s). Effect of caffeine solution stimulation (10mM). GFP signal was recorded every 500 ms.

**Extended data Movie 3 | Gr66a+ neurons respond *in vivo* to PGN.**

Real-time calcium imaging using the calcium indicator GCaMP6s to assess the *in vivo* neuronal activity in the sub-esophageal zone of bitter neurons (Gr66a^Gal4^/UAS-GCaMP6s). Effect of peptidoglycan solution stimulation (100 μg/mL). GFP signal was recorded every 500 ms.

## Methods

### Fly stocks

All flies were maintained at 25°C on a standard cornmeal/agar medium on a 12 h:12 h light-dark cycle with a relative humidity of 70%.The strains used are the following: pLB1^Gal4 15^; PGRP-LB::GFP ^16^; w (BDSC:3605); yw; Canton-S; Gr5a^LexA 44, 45^ (Gently provided by Dong Min Shin); Gr66a^LexA^ (^46^; gently provided by K. Scott’s Lab); Gr32a^LexA 47^ (gently provided by A. Dahanukar’s lab); Gr32a^Gal4^ (BDSC:57622); Gr66a^Gal4^; Gr66a-RFP(X4) (BDSC:60691); UAS-TrpA1 (BDSC:26264,^48^); UAS-Kir2.1 (BDSC:6595); 40XUAS-mCD8-GFP (BDSC:32195); UAS-Fadd RNAi^49^; UAS-Imd RNAi (VDRC#101834); UAS-Dredd-RNAi (VDRC#104726); UAS-PGRP-LC RNAi (VDRC#101636); UAS-Relish RNAi (BDSC:28943); UAS*frt*STOP*frt* mCD8GFP (BDSC:30125); 8XLexAop2-FLP (BDSC:55819); UAS-GCaMP6s (BDSC:42746). UAS-PGRP-LCa ^50^; PGRP-LC^E12 51^; PGRP-LE^112 52^, Dredd^D55 53^; TrpA1^1 21^.

### Tastants

For *in vivo* calcium imaging and flyPAD assays tastants were dissolved in autoclaved purified distilled water. All tastant solutions were freshly prepared and stored in aliquots at −20°C for a maximum duration of six months. Peptidoglycan was obtained from InvivoGen (PGN-EK Catalog # tlrl-pgnek, InvivoGen, USA), while sucrose (Roth, ref 4621.1) and caffeine (Sigma Aldrich, ref C0750) were obtained from Sigma-Aldrich (USA).

### *In vivo* calcium imaging

*In vivo* calcium imaging experiments were performed on 5-7-day-old mated females. Flies were starved for 24 h before any experiments. Flies of the appropriate genotype were anesthetized on ice for 1 h. Female flies were suspended by the neck on a plexiglass block (2 × 2 × 2.5 cm), with the proboscis facing the center of the block. Flies were immobilized using an insect pin (0.1 mm diameter) placed on the neck. The ends of the pin were fixed on the block with beeswax (Deiberit 502, Siladent, 209212). The head was then glued on the block with a drop of rosin (Gum rosin, Sigma-Aldrich −60895-, dissolved in ethanol at 70 %) to avoid any movements. The anterior part of the head was thus oriented towards the objective of the microscope. Flies were placed in a humidified box for 1 h to allow the rosin to harden without damaging the living tissues. A plastic coverslip with a hole corresponding to the width of the space between the two eyes was placed on top of the head and fixed on the block with beeswax. The plastic coverslip was sealed on the cuticle with two-component silicon (Kwik-Sil, World Precision Instruments) leaving the proboscis exposed to the air. Ringer’s saline (130 mM NaCl, 5 mM KCl, 2 mM MgCl_2_, 2 mM CaCl_2_, 36 mM saccharose, 5 mM HEPES, pH 7.3, was placed on the head^54^. The antenna area, the tracheas, and the fat body were removed. The gut was cut without damaging the brain and taste nerves to allow visual access to the anterior ventral part of the sub-esophageal zone. The exposed brain was rinsed twice with Ringer’s saline. GCaMP6s fluorescence was viewed with a Leica DM600B microscope under a 25x water objective. GCaMP6s was excited using a Lumencor diode light source at 482 nm ± 25. Emitted light was collected through a 505-530 nm band-pass filter. Images were collected every 500 ms using a Hamamatsu/HPF-ORCA Flash 4.0 camera and processed using Leica MM AF 2.2.9. Stimulation was performed by applying 140 μL of tastant solution diluted in water on the proboscis. Each experiment consisted of a recording of 10 images before stimulation and 30 images after stimulation. Data were analyzed as previously described by using FIJI (https://fiji.sc/)^54^. In all experiments implicating pLB1^Gal4^, this driver and the UAS-GCaMP6s transgenes are homozygous. In experiments using Gr66a^Gal4^, the driver and the UAS-GCaMP6s transgenes are heterozygous.

### Immunostaining and imaging

Immunostaining and imaging were performed as previously described ^16^. Brains from adult females were dissected in Phosphate-buffered saline (PBS, Eurobio, ref CS0PBS0108) and fixed for 15 min in 4% paraformaldehyde (Electron Microscopy Sciences, Cat # 15714-S) at room temperature (RT). Afterward, brains were washed three times for 10 min in PBS-T (PBS + 0.3% Triton X-100) and blocked in 2,5% bovine serum albumin (BSA; Sigma-Aldrich) in PBS-T for 30 min. After saturation, samples were incubated with the first antibody diluted in 0,5% BSA in PBS-T overnight at 4°C. The following day, brains were washed three times and incubated with the secondary antibody diluted in 0,5% BSA in PBS-T for 2h at RT. Next, samples were washed for 10 min in PBS-T and mounted on slides using Vectashield (Vector Laboratories, Ca, USA) fluorescent mounting medium. In the case of proboscises, no immunostaining was performed. Proboscises of adult females were dissected in PBS, rinsed with PBS and directly mounted on slides using Vectashield fluorescent mounting medium. The tissues were visualized directly after.

For the immunostaining the first antibodies used are the following: Chicken anti-GFP (Aves Labs Cat#GFP-1020, RRID:AB_10000240. Dilution 1:1000), rabbit anti-RFP (Rockland Cat#600-401-379, RRID:AB_2209751. Dilution 1:1000), mouse anti-NC82 (DSHB Cat#nc82,RRID:AB_2314866. Dilution 1:40). The secondary antibodies used are the following: Alexa Fluor 488 Donkey anti-Chicken IgY (IgG) (H+L) (Jackson ImmunoResearch Labs Cat#703-545-155, RRID:AB_2340375. Dilution 1:500), Alexa Fluor568 donkey anti-mouse IgG (H+L) (Thermo Fisher Scientific Cat#A10037, RRID:AB_2534013. Dilution 1:500), Alexa Fluor647 donkey anti-mouse IgG (H+L) (Jackson ImmunoResearch Labs Cat#715-605-151, RRID:AB_2340863. Dilution 1:500), Alexa Fluor 568 donkey anti-rabbit IgG (H+L) (Thermo Fisher Scientific Cat#A10042, RRID: AB2534017. Dilution 1:500).

Images were captured with either a Leica SP8 confocal microscope (in this case, tissues were scanned with 20X oil immersion objective) or an LSM 780 Zeiss confocal microscope (20x air objective was used). For the detection of endogenous PGRP-LB::GFP, images were captured with a Spinning Disk Ropper 2 Cam (20x or 40x air objective were used). Images were processed using Adobe Photoshop.

### Feeding assay

Two-choice feeding assays were performed by using the flyPAD device ^32^ which records the cumulative number of sips. Each sip corresponds to a contact of flies’ proboscis with the chosen food substrate. Individual 5–7-day-old flies were starved for around 20h and then aspirated in arenas containing two food substrates. The control substrate consisted in a 1% agarose 5mM sucrose solution, whereas the test substrate additionally contained peptidoglycan at the indicated concentrations. Each arena’s well (2 per arena) was filled with 3.5 μL of food solution. Tests were ran for 1h at 25 °C. Data were collected and analyzed by using Bonsai ^55^and MATLAB, respectively (scripts provided by Pavel Itskov). Preference index was calculated as following: (number of sips in the test solution - number of sips in the control solution)/ total number of sips. Non eaters were excluded from the analysis.

### Oviposition assay

Oviposition assays were performed as previously described ^16^. Eclosed flies were raised at 25°C or RT, in case of experiments involving the thermosensitive transgene UAS-TrpA1. 5-day-old mated females were anesthetized on a CO_2_ pad and singularly transferred in tubes with fresh media and dry yeast (Fermipan) added on top of each tube right before the egg-lay period. Flies were let to lay eggs for 24 h at 29°C or 23°C in control conditions for experiments involving UAS-TrpA1. After the 24 h, eggs were counted.

### Statistical analysis

GraphPad Prism software was used for statistical analyses. For *in vivo* Calcium imaging and feeding assay analysis non-parametric unpaired Mann-Whitney two-tailed tests were performed. In the case of oviposition assay, we used the non-parametric unpaired ANOVA, Kruskal-Wallis test, and Dunn’s post-test.

## Acknowledgments

We thank Emilie Avazeri and Annelise Viallat-Lieutaud for technical help. We thank members of the Royet’s laboratory for their comments on the manuscript. This work was supported by (ANR-11-LABX-0054) (Investissements d’Avenir–Labex INFORM), ANR BACNEURODRO (ANR-17-CE16-0023-01) and ANR PEPTIMET (ANR-18-CE15-0018-02), Equipe Fondation pour la Recherche Médicale (EQU201903007783) and l’Institut Universitaire de France to J.R. Y.G. laboratory is supported by the “Centre National de la Recherche Scientifique”, the “Université de Bourgogne Franche-Comté”, the Conseil Régional Bourgogne Franche-Comte (PARI grant), the FEDER (European Funding for Regional Economical Development), and the European Council (ERC starting grant, GliSFCo-311403).

## Author contributions

Genetic epistasis and imaging and behavioral assay were performed by A.M. and L.K. Calcium imaging was performed by G.M. Results were analyzed and interpreted by A.M., G.M., Y. G., L. K., and J. R. The original draft was written by J.R. Reviewing and editing were performed by all authors. Supervision: L.K., Y.G., and J.R. Funding acquisition: Y.G, and J.R.

